# Non-invasive muscle biopsy: estimation of muscle fibre size from a neuromuscular interface

**DOI:** 10.1101/2022.10.21.513157

**Authors:** Andrea Casolo, Sumiaki Maeo, Thomas G. Balshaw, Marcel B. Lanza, Neil R. W. Martin, Stefano Nuccio, Tatiana Moro, Antonio Paoli, Francesco Felici, Nicola Maffulli, Bjoern Eskofier, Thomas M. Kinfe, Jonathan P. Folland, Dario Farina, Alessandro Del Vecchio

**Affiliations:** Department of Biomedical Sciences, University of Padova, Padua, Italy; Faculty of Sport and Health Science, Ritsumeikan University, Kusatsu, Shiga, Japan; School of Sport, Exercise & Health Sciences, Loughborough University, United Kingdom; Versus Arthritis Centre for Sport, Exercise and Osteoarthritis Research, Loughborough University, Leicestershire, United Kingdom; Department of Physical Therapy and Rehabilitation Science, University of Maryland, Baltimore, Maryland; Department of Movement, Human and Health Sciences, University of Rome ‘Foro Italico’, Rome, Italy; Department of Musculoskeletal Disorders, University of Salerno, Salerno, Italy; Queen Mary University of London, Barts and the London School of Medicine and Dentistry, Centre for Sports and Exercise Medicine, Mile End Hospital, London, United Kingdom; Department Artificial Intelligence in Biomedical Engineering, Friedrich-Alexander University, Erlangen-Nürnberg, 91052, Erlangen, Germany; Department of Bioengineering, Imperial College London, London, United Kingdom

**Keywords:** Motor unit, muscle fibre area, conduction velocity, high-density surface electromyography.

## Abstract

Because of the biophysical relation between muscle fibre diameter and the propagation velocity of action potentials along the muscle fibres, motor unit conduction velocity (MUCV) could be a non-invasive index of muscle fibre size in humans. However, the relation between MUCV and fibre size has been only assessed indirectly in animal models and in human patients with invasive intramuscular EMG recordings, or it has been mathematically derived from computer simulations. By combining advanced non-invasive techniques to record motor unit activity in vivo, i.e., high-density surface EMG, with the gold standard technique for muscle tissue sampling, i.e., muscle biopsy, here we investigated the relation between the conduction velocity of populations of motor units identified from the biceps brachii muscle, and muscle fibre diameter. Moreover, we demonstrate the possibility to predict muscle fibre diameter (R^2^ = 0.66) and cross-sectional area (R^2^ = 0.65) from conduction velocity estimates with low systematic bias (~2% and ~4% respectively) and a relatively low margin of individual error (~8% and ~16%, respectively). The proposed neuromuscular interface opens new perspectives in the use of high-density EMG as a non-invasive tool to estimate muscle fibre size without the need of surgical biopsy sampling. The non-invasive nature of high-density surface EMG for the assessment of muscle fibre size may be useful in studies monitoring child development, aging, space and exercise physiology.

**SIGNIFICANCE STATEMENT:** Our study explored the relation between the conduction velocity of populations of motor units and muscle fibre size in healthy humans. Our results provide in vivo evidence that a high-density surface EMG-derived physiological parameter, i.e. motor unit conduction velocity, can be adopted to estimate muscle fibre size, without the need of surgical biopsy sampling. Here we propose a neuromuscular interface that opens new perspectives not only in the study of neuromuscular disorders, but also in other fields where the non-invasive and painless determination of muscle fibre and motor unit size becomes a priority, such as in aging, space and exercise physiology.

## INTRODUCTION

The motor unit is the basic functional element of the neuromuscular system, whose primary function is to transduce the neural activation signal into muscular forces and movement (1). It consists of a motor neuron and the group of muscle fibres it innervates (1, 2). Skeletal muscles are composed of muscle fibres, whose heterogeneity results from the broad range of properties of both the innervating motor neuron and the structural and biochemical composition of the fibres (2, 3).

Since the pioneering work of Adrian and Bronk (4), different techniques have been developed to record motor unit activity in humans either invasively or non-invasively (5). Although intramuscular EMG recordings remain the classic means for investigating the activity and properties of individual motor units, advances in electrode fabrication, biosignal processing and modeling over the last twenty years, have contributed to expand the opportunities to study motor unit activity in humans non-invasively (6). For instance, the recent introduction of high-density surface EMG (HDsEMG) have greatly helped to overcome some of the limitations of intramuscular EMG recordings (6–8), making it possible to decode and monitor the concurrent activity of large and representative populations of motor units non-invasively (9, 10). These arrays of electrodes enable the sampling of individual motor unit action potentials (MUAPs) with high spatial and temporal accuracy (11, 12).

On the other hand, the muscle biopsy technique provides an invaluable opportunity to study several characteristics of skeletal muscle tissue and muscle fibre properties, including fibre type composition and cross-sectional area (CSA) (13, 14). In turn, the study of these features may yield essential information about muscle function, dysfunction and adaptability (15, 16). In this respect, the seminal work by Charriere and Duchenne in 1865 (17), which described the first known percutaneous biopsy needle (i.e., *emporte-pièce histologique*), heralded a new era in the study of a wide range of health and disease conditions. Since then, methodological advances in biopsy sampling and processing of the specimen, have contributed to spread the use of the muscle biopsy procedure beyond its original context of clinical diagnosis of neuromuscular disorders, to become the gold standard technique for the evaluation of fibre morphology within general physiology and exercise sciences (18, 19).

For instance, the use of muscle biopsy has been fundamental for our understanding of common morphological changes in response to aging, disease, disuse/unloading (e.g. disuse, immobilization) or use/loading (20, 21). In this respect, the increased muscle fibre CSA, i.e., fibre hypertrophy, the preferential hypertrophy of type II fibres, as well as the subtle modifications in fibre type and myosin heavy-chain composition, are well-documented adaptations underlying the enhancements in volitional force production after prolonged mechanical loading (e.g. resistance training).

However, although safe and with a low rate of complications, the biopsy method requires the insertion of the needle through the skin and muscle fascia, which exposes the individual to some risks both during and after the procedure (18, 22). The poor reproducibility of this technique for fibre area measurements (19, 23–25), and the fact that it provides morphological information dissociated from the neural control of the sampled muscle fibres (26), have also been well-documented. Thus, it would be enticing to identify alternative physiological parameter, which could inform us about the size of muscle fibres belonging to individual motor units non-invasively.

One potential candidate to indirectly estimate fibre size is muscle fibre conduction velocity (MFCV), which can be measured from surface EMG. The inverse of the average latency between surface EMG signals recorded from adjacent electrodes aligned in the direction of the muscle fibres, provides an estimate of the average propagation velocity of the MUAPs of several concurrently active motor units (27). Alternatively, by decomposing the surface EMG signal and separating MUAPs of individual motor units (28, 29), it is possible to estimate the motor unit conduction velocity (MUCV) (30, 31). Although both parameters can be estimated non-invasively from surface EMG, a major advantage of MUCV over MFCV estimates is that the former provides information about individual motor unit properties and not the average properties of all active motor units.

MUCV is considered a “size principle parameter”, due to its linear association with motor unit recruitment threshold (26, 32, 33), thus it has been generally adopted to indirectly infer neural control strategies (34–36). This linear association is due to the direct relation between muscle fibre diameter (MFD) and conduction velocity (37, 38), observed with invasive techniques in animal models (37) and humans (38, 39). Therefore, the possibility of estimating the conduction velocity for a large population of MUs non-invasively would theoretically allow an estimate of the distribution of muscle fibre sizes alternative to muscle biopsies.

By combining HDsEMG (11) with the gold standard technique for muscle tissue sampling (i.e., muscle biopsy), here we first systematically investigated the relation between the conduction velocity of large populations of motor units identified non-invasively during voluntary contractions, and muscle fibre size, i.e., MFD and fibre CSA, directly derived from muscle biopsies. We studied this relation in the biceps brachii (BB) muscle of a heterogeneous population consisting of healthy untrained and chronically resistance-trained subjects, in order to have a sample of widely varying muscle fibre size. From this analysis, we then propose a new methodology to estimate the average MFD and fibre CSA from surface EMG processing without the need for muscle biopsies.

Thus, the main purpose of this study was to determine how MUCV and muscle fibre size (MFD and fibre CSA) were related in an heterogenous population of healthy adults, by considering inter-participant variability of both parameters. We hypothesized that MUCV, assessed in the present study from large populations of concurrently active motor units, would have a strong association with MFD, thus becoming a novel non-invasive marker of fibre size. We reveal that it is possible to accurately transform MUCV values into a distribution of fibre diameters, therefore opening new avenues for research in ageing, training, and neuromuscular disease.

## RESULTS

We report a systematic assessment of the relation between MUAPs propagation velocity of individual motor units, i.e., MUCV, here estimated non-invasively from large and representative populations of motor units during isometric voluntary contractions, and the size of muscle fibres, i.e., MFD and fibre CSA, directly derived from muscle biopsies, in an heterogeneous population of healthy individuals. Figure 1 shows an overview of the study.

**Figure 1.**
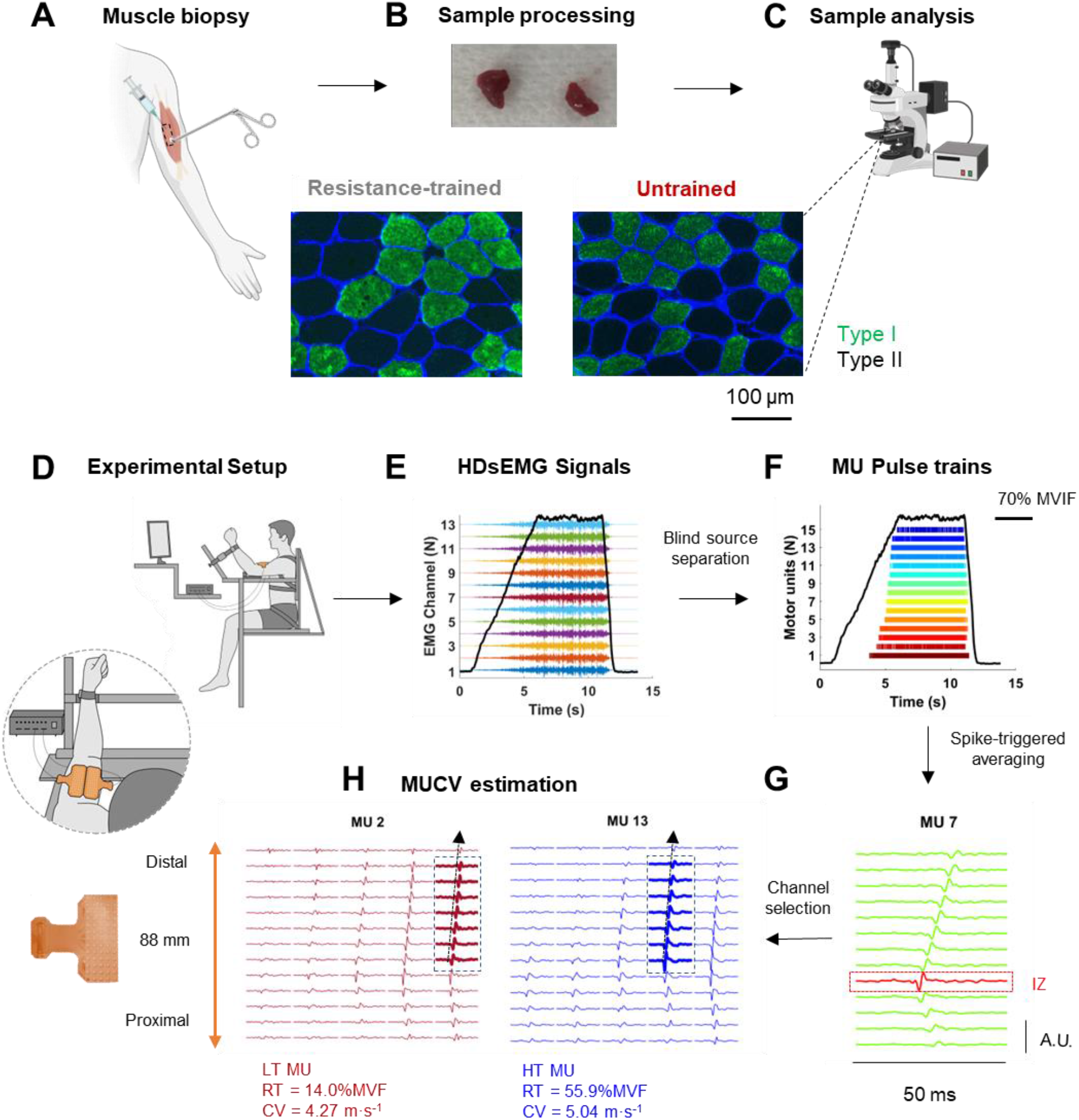
(A) Muscle biopsy was taken from the biceps brachii (BB) muscle from the same region where HDsEMG recordings were made. (B) Transverse serial cross sections (8 μm thick) were obtained using a cryotome before being incubated with a primary antibody for myosin heavy chain I. (C) Images were captured using a fluorescence microscope. Fibre cross-sectional area (fibre CSA) was assessed by manually drawing around the perimeter of each fibre for ~200 different fibres per participant. Representative examples of type I and II fibre CSA measured from one resistance-trained (RT) and one untrained control (UT) participant are displayed. (D) Experimental configuration for the main neuromuscular assessment. Two HDsEMG grids were placed over the BB muscle to cover most of the skin surface overlying the muscle. (E) Representative example of a linearly increasing isometric ramp contraction to 70%MVF (force signal in black), with concomitant monopolar recordings from 13 electrodes from one column of the grid (displayed in different colors). (F) Color-coded raster plots of the individual motor units (n = 15) identified from BB muscle for one representative participant. (G) Motor unit action potential (MUAPs) waveforms from the individual pulse trains of the identified motor units were obtained via the spike triggered averaging technique. Clear propagation of a single MUAP from the innervation zone (IZ, in red) both to the proximal and the distal tendon regions along one electrode column can be observed. (H) Motor unit conduction velocity (MUCV) was estimated with a validated multichannel maximum likelihood algorithm. A representative example of a low threshold motor unit (LT MU, in red) and of a high threshold motor unit (HT MU, in blue) is shown. The MUAP of the HT MU (5.04 m·s^−1^) propagates with a greater velocity than that of the LT MU (4.27 m·s^−1^).

### High-density EMG decomposition and motor unit properties

Myoelectrical signals recorded during voluntary ramp contractions were decomposed into discharge instants of individual motor units (12, 28) (Figure 1 D-F). After visual inspection and manual editing (see *HDsEMG Decomposition* in Methods), when considering all the 29 participants and the four contraction levels performed by each participant, we identified a total of 701 unique motor units within the 29 BB short-head muscles. The average number of identified motor units per participant was 24.2 ± 8.6 (range 11 to 47).

The average normalized MU RT, i.e., the force value relative to maximum (%MVF) at which the first MUAP was discharged, was 31.5 ± 4.0 %MVF (range 24 to 42.7). Based on their normalized MU RT, the identified motor units were clustered into low-threshold (LT MU, range of MU RT 0-30 %MVF) and high-threshold (HT MU, range of MU RT 50-70 %MVF) (26, 40). A total of 298 and 76 motor units were classified as LT and HT MUs, respectively. The average number of identified LT and HT MUs per participant was 10.3 ± 4.3 (range 2 to 22) and 2.6 ± 2.2 (range 1 to 9), respectively.

For each participant, the average recruitment MU DR, i.e., the mean of the instantaneous discharge rates of the first 20 MUAPs in the ascending phase of the ramp contractions, was 17.4 ± 2.7 pulses per second (pps) (range 14.3 to 23.2 pps).

### Motor unit conduction velocity

The conduction velocity of the identified motor units was estimated with a validated multi-channel maximum likelihood algorithm (see *Motor unit conduction velocity estimation* in Methods). Figure 1G shows a representative example of one motor unit with MUAPs propagating from the innervation zone (in red) in both directions along an electrode column of the grid.

Figure 2B shows the relative frequency histogram for MUCV values of all the identified motor units (n = 701; range 3.77 to 6.02 m·s^−1^). In agreement with previous observations (26, 41), the distribution was unimodal and did not highlight distinct clusters of MUCV values. The average conduction velocity (MUCV_AVERAGE_) of each participant from all their active motor units in the ascending phase of the ramp contraction, was 4.76 ± 0.33 m·s^−1^ (range 4.34 to 5.48 m·s^−1^).

**Figure 2.**
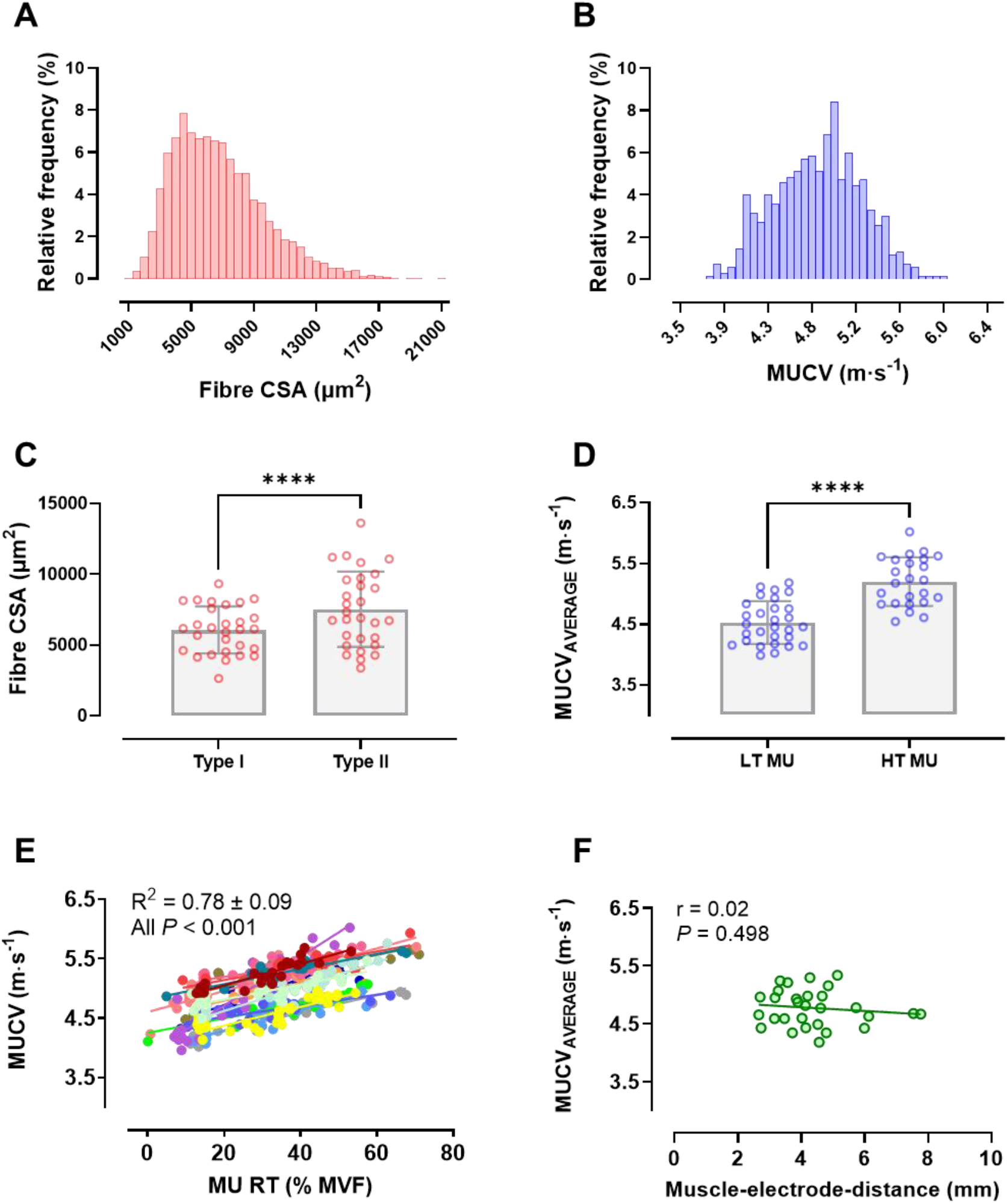
Relative frequency histogram of (A) fibre cross sectional area (Fibre CSA) of all the analyzed muscle fibres (n = 5619; range 1127 to 20803 μm^2^); and (B) motor unit conduction velocity (MUCV) of all identified motor units (n = 701; range 3.77 to 6.02 m·s-1). (C) Bar plots displaying the average type I and type II fibre CSA measured from muscle biopsies for each participant. For each participant, average type II fibre CSA was significantly greater than that of type I fibres (*P* = 0.015). (D) Bar plot displaying the average motor unit conduction velocity (MUCV_AVERAGE_) of low threshold (LT MU) and high threshold (HT MU) for each participant. The average propagation velocity of HT MU was significantly higher than that of LT MU (*P* < 0.001). (E) MUCV regression lines plotted as a function of normalized motor unit recruitment threshold (MU RT) for sixteen representative participants. Each dot represents a single MUCV estimate and each color represent a different participant. MUCV linearly increased as a function of normalized MU RT in all participants. The average coefficient of determination (R^2^) across participants is reported in the upper left corner. (F) Correlation analysis between average muscle-electrode-distance (MED) and MUCV_AVERAGE_ of each of the 29 participants. Each grey dot represents a different participant. The coefficient of correlation (r) is displayed in the upper left corner.

HT MU displayed significantly higher MUCV compared to that of LT MU (5.20 ± 0.40 m·s^−1^ vs. 4.53 ± 0.35 m·s^−1^, *P* < 0.001, Figure 2 D). As expected, the faster propagation velocity of MUAP in HT MU compared to LT MU was further confirmed by the significant linear relation between MUCV and MU RT, which was observed in each of the 29 participants. Individual participant R^2^ values for the relationship of MUCV and MU RT ranged from 0.59 to 0.93 (*P* < 0.001, in all participants) with an average of 0.78 ± 0.09. Representative examples of the relation between MUCV and MU RT from 16 participants are shown in Figure 2 E.

In order to assess the influence of subcutaneous adipose tissue thickness (i.e., volume conductor) on MUCV estimates, we also assessed the inter-individual association (i.e., n = 29) between the mean muscle-electrode-distance (MED), measured with B-mode ultrasonography from the BB short head (see *Ultrasound recording* in Methods), and MUCV_AVERAGE_. The absence of a significant correlation between the two parameters (*P* = 0.498, Figure 2 F) indicated that MUCV values estimated from surface HDsEMG decomposition were not influenced by the subcutaneous adipose tissue interposed between the muscle belly and the recording site. This rules out the possibility that adipose tissue would be a confounding factor for the results.

### A non-invasive interface to estimate muscle fibre size

Single MUCV values were converted into estimated MFD and subsequently related to the measured MFD directly obtained from muscle biopsies. By relating the estimated MFD with actual measured MFD, we developed a non-invasive interface which allowed the prediction of the mean measured MFD and of fibre CSA at the individual subject level from an EMG-derived neuromuscular parameter (see *Estimation of muscle fibre diameters from motor unit conduction velocity* in Methods).

Figure 3A shows the relative frequency histograms of measured and estimated MFD, when pooling all analyzed muscle fibres (n = 5619, in red) from all participants. Measured MFD ranged from 37.9 to 162.8 μm, with an average value of 90.8 ± 19.6 μm. Similarly, when considering all identified motor units (n = 701, in blue), estimated MFD ranged from 41.6 to 148.7 μm, with an average value of 90.8 ± 19.7 μm, following a calibration (see Methods). The estimated MFD distribution was very similar to that of the measured MFD, which showed that the calibration procedure allowed to almost perfectly map MUCV into MFD values. The similarity between the measured and estimated MFD distributions was further confirmed by the Kolmogorov-Smirnov test (D = 0.050, *P* = 0.086). It has to be noted that, while the agreement of mean and standard deviation of the two distributions was expected because of the calibration, the non-significant difference between the entire distributions of MUCV and MFD indicated similar shapes of the two histograms that could not be due only to the calibration (which was a linear transformation).

**Figure 3.**
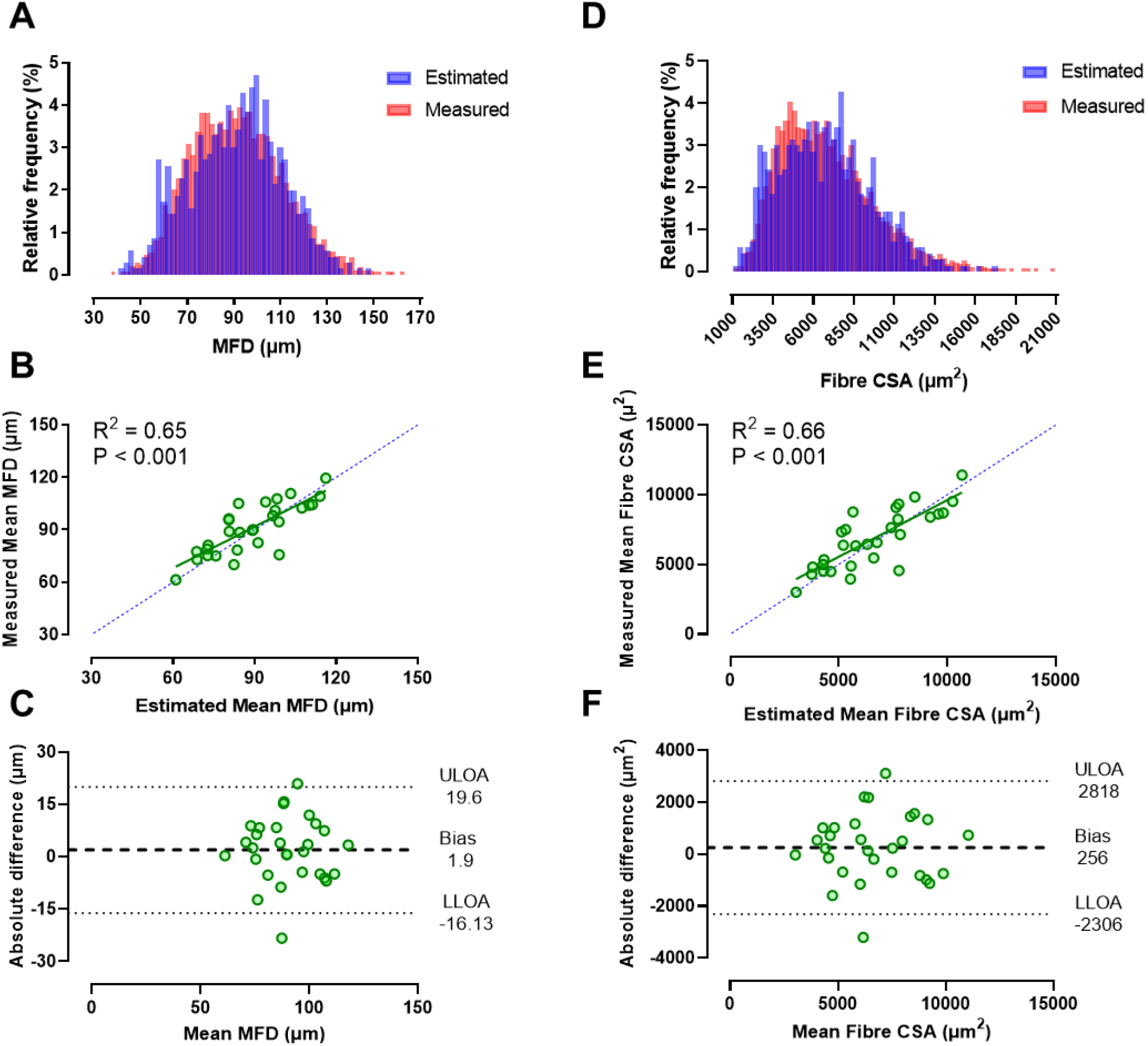
Relative frequency histograms of (A) muscle fibre diameters (MFD); and (D) muscle fibre cross-sectional area (fibre CSA), when considering all individual data from the 29 participants. In both graphs, the distributions of measured MFD (A) and measured fibre CSA (D), directly derived from muscle biopsies, are shown in red, whereas the distributions of estimated MFD (A) and estimated fibre CSA (D), derived from motor unit conduction velocity, are shown in blue. Measured and estimated MFD distributions (A) were composed of 170 bins with a ~2 μm width; Measured and estimated fibre CSA distributions (D) were composed of 81 bins with a ~250 μm^2^ width. Inter-subject relations between (B) estimated mean MFD and measured mean MFD; and (E) estimated mean fibre CSA and measured mean fibre CSA. Linear regression analyses confirmed that (B) the estimated mean MFD can predict the majority of the variance of the measured mean MFD, at the individual subject level. Similarly, (E) the estimated mean fibre CSA can predict the majority of the variance of the measured mean fibre CSA, at the individual subject level. The coefficient of determination (R^2^) is displayed in the upper left corner. Each green dot represents a different participant. Bland-Altman analyses showing absolute differences between (C) estimated mean MFD and measured mean MFD; and (F) between estimated mean fibre CSA and measured mean fibre CSA in absolute values. A mean absolute bias in (C) estimated mean MFD compared with measured mean MFD of 1.9 ± 9.2 and in (F) estimated mean fibre CSA compared with measured mean fibre CSA of - 256 ± 1307 μm^2^ were detected. ULOA, upper limit of agreement; LLOA, lower limit of agreement.

The distributions of MUCV and MFD obtained from the entire dataset provide an indication of the relation between propagation velocity and fibre diameter in our dataset. *Equation 3* in Methods provides the linear transformation to obtain conduction velocity from fibre diameter and vice versa, according to the estimates of conduction velocity obtained by the proposed technique. The next step was to test whether the relation between the two variables could only be obtained for the full population or whether it was robust enough to allow for the prediction of the distribution of fibre MFD given the measured distribution of MUCV for individual subjects. Basically, we wanted to study the predictability of MFD distribution for each subject when the MUCV was measured. For this analysis, we obtained the linear relation between MUCV and MFD for each subject by a leave-one-out procedure. According to this procedure, the optimal fitting parameters for *Equation 2* (see Methods) were obtained using the entire dataset but excluding the subject for whom the estimates were derived. Once the parameters were determined using data from all subjects but one, the MFD distribution for the left-out subject was estimated using *Equation 2* with the optimal parameters obtained from the remaining dataset. This was repeated by leaving out all subjects, one at a time, so that estimated MFD distributions were obtained for each subject without using the corresponding subject-specific biopsy data. This procedure corresponded to estimating MFD distributions from a general association with MUCV obtained from a training/calibration set of data. After this operation, we compared the measured MFD distributions from each subject with those estimated from the MUCV distributions to which the subject-specific linear transformation derived from all other subjects was applied. In this way, the calibration was obtained from data not used for the analysis of the association between estimated and measured MFD distributions at the individual subject level. This approach provided information on the predictability of MFD at the individual subject level, following a separate calibration on a representative sample of individuals. With this procedure, the Pearson’s correlation analysis revealed a strong and significant correlation between measured and estimated mean MFD across individuals (r = 0.81, 95% CI = 0.62 0.90, *P* < 0.001). Regression analysis further confirmed that estimated mean MFD was able to predict the majority of the variance of the measured mean MFD (R^2^ = 0.65, *P* < 0.001, Figure 3 B). This indicated that the MFD could be estimated using only the MUCV data with a calibration on a separate dataset. For new subjects, it would be therefore possible to estimate MFD distributions without needing the biopsy data. According to high level of predictability obtained on an individual subject basis, the parameters estimated for each subject with the leave-one-out procedure varied in a very small range when changing the left-out subject.

Bland-Altman plots are shown in Figure 3 C for absolute values and revealed a mean absolute bias in estimated mean MFD compared with measured mean MFD of 1.9 ± 9.2 μm across all subjects. The mean relative bias across subjects was 2.2 ± 10.4%. Moreover, the mean absolute and relative errors in estimated mean MFD compared with measured mean MFD across subjects were 7.3 ± 5.8 μm and 8.1 ± 6.8%, respectively.

Lastly, considering that fibre CSA is generally regarded as a more direct and precise measure of fibre size in humans compared to MFD, we also explored the relation between muscle fibre CSA estimated from MUCV and the muscle fibre CSA directly measured from muscle biopsies. Firstly, estimated MFD data from all participants were converted into estimated fibre CSA (see *Equation 4* in Methods). Figure 3 D shows the relative frequency histograms of measured and estimated fibre CSA, when pooling individual data from all participants (n = 701). Similarly to what was observed for MFD, the estimated fibre CSA distribution (6782.0 ± 2829.1 μ^2^) was very similar to that of the measured fibre CSA (6771.8 ± 2912.0 μ^2^), as shown by the almost identical means and standard deviations. The similarity between the measured and estimated CSA distributions was further confirmed by the Kolmogorov-Smirnov test (D = 0.049, *P* = 0.090). As for MFD, this was expected because of the global calibration. Moreover, at the individual subject level, the regression analysis revealed a strong and significant association between measured and estimated mean fibre CSA (R^2^ = 0.66, *P* < 0.001, Figure 3 E) (this was also obtained with the leave-one-out procedure, as discussed above).

In contrast to the findings for fibre size (MFD and fibre CSA) and MUCV, there was no association between MUCV and muscle fibre type composition (i.e. %type II fibres by area; r = 0.19, *P* = 0.302), as well as between fibre CSA and muscle fibre type composition (i.e. %type II fibres by area; r = 0.30, *P* = 0.105).

Bland-Altman plots are shown in Figure 3 F for absolute values and indicate a mean absolute bias in estimated mean fibre CSA compared with measured mean fibre CSA of 256 ± 1307 μm^2^ across subjects. The mean relative bias across subjects was 4.0 ± 20.3%. Moreover, the mean absolute and relative errors in estimated mean fibre CSA compared with measured mean fibre CSA across subjects were 1037 ± 814 μm^2^ and 15.9 ± 14.4%, respectively.

These results suggest a good degree of accuracy in predicting important indicators of muscle fibre size, i.e., the mean MFD and mean fibre CSA, with our non-invasive neuromuscular interface.

## DISCUSSION

We demonstrated that a non-invasive neuromuscular interface allows the prediction of the mean muscle fibre size (i.e., MFD and fibre CSA) for a heterogeneous population of healthy individuals with a relatively low margin of individual error (≤16%) and very low systematic bias (≤4%). The strong and positive relations observed between the MFD/fibre CSA estimated from MUCV and the MFD/fibre CSA directly derived from muscle biopsies (R^2^ = 0.65 and R^2^ = 0.66, respectively), show experimentally for the first time that MUCV may be adopted as a novel strong index of muscle fibre size in healthy individuals. Our results provide substantial novelty about the potential adoption of an EMG-derived, and fully non-invasive, physiological parameter to estimate muscle fibre size with a relatively low margin of error.

We combined the state-of-the-art technique (12, 28) for the non-invasive recording of representative populations of concurrently active motor units (i.e., HDsEMG), with the gold-standard technique for the assessment of fibre size and composition in humans (i.e., muscle biopsy). Through HDsEMG, we decoded the activity and extracted the properties (i.e., MU RT, MU DR and MUCV) of 701 unique motor units voluntarily activated in the BB muscle during isometric elbow flexions. The accuracy and validity of the decomposition technique for motor unit identification adopted in this study have been tested in a number of previous reports (11). Similarly, through muscle biopsy we directly measured the size of 5619 individual muscle fibres from the same region of BB muscle from which HDsEMG was recorded.

The main finding of the current analysis is that we revealed the possibility to transform MUCV data into estimated measures of muscle fibre size (i.e., estimated MFD and fibre CSA), which in turn showed a good degree of association with the actual muscle fibre size measured directly by muscle biopsy. Firstly, this was done by a calibration across the full subject sample that identified the two parameters of Nandedkar and Stålberg’s equation (*Equation 2*) that minimized the distance between estimated and measured MFD. The two optimized parameter values (*Equation 3*) corresponded to the best fit between MUCV and MFD, as directly derived from muscle biopsies (Figure 3 A; Kolmogorov-Smirnov test: D = 0.050, *P* = 0.086). Secondly, the proposed best fitting linear equation was used to transform MUCV estimates into MFD in order to assess the relation between estimated mean MFD and measured mean MFD in individual subjects. Regression analysis between the measured and estimated mean MFD demonstrated the possibility to predict the mean measured MFD (Figure 3 B, R^2^=0.65) from an EMG-derived parameter across subjects, with a relatively low bias (Figure 3 C; mean absolute bias: 1.9 μm; mean relative bias: 2.2%) and error (mean absolute error: 7.3 μm; mean relative error: 8.1%).

Since muscle fibre CSA is generally considered as a more precise indicator of fibre size, because of the typical irregular shape of skeletal muscle fibres, we also assessed if the proposed model allowed an accurate prediction of measured fibre CSA. Our results confirmed that it was possible to predict the mean measured fibre CSA (Figure 3 E) from MUCV across subjects, with a relatively low bias (Figure 3 F; mean absolute bias: 256 μm^2^; mean relative bias: 4.0%) and error (mean absolute error: 1037 μm^2^; mean relative error: 15.9%).

The proposed equation which describes the linear relation between MUCV and MFD (*Equation 3*) presents normalization values (constants) that are somewhat different to those observed by previous studies on both healthy (42) and pathological (38) individuals. These discrepancies may be due to several methodological differences. First, in these previous studies the conduction velocity has been estimated with intramuscular EMG (e.g., concentric needle electrode), during electrically-elicited stimulations in resting conditions (38) or it has been mathematically-derived from simulations of intramuscular recordings (42). In the first case, in addition to the invasiveness of the procedure used for its estimation, the high selectivity associated with intramuscular EMG recordings combined with electrical stimulations may have hindered the sampling of motor units in a representative way, a condition which may have affected the apparent relations between the conduction velocity and MFD (38). Moreover, the estimates of conduction velocity from intramuscular EMG are generally lower, in the order of ~1.0 m·s^−1^, and more variable than the estimates obtained from the surface EMG (43). In the second case, the adoption of simulated intramuscular EMG recordings for conduction velocity estimation did not allow a systematic assessment of the relation with MFD based on intrasubject observations of both measurements from the same muscle (42), as was done in the current analysis.

Conversely, in the current study we derived the MUCV of large and representative populations of motor units concurrently active during a voluntary task in a fully non-invasive way and we systematically assessed its relation with large samples of biopsy-derived MFD. It is worth noting also, that contrary to previous studies that directly (38, 39) or indirectly (42) assessed the relation between conduction velocity and muscle fibre size, current MUCV estimates were obtained with the most accurate and validated algorithm available, which ensured a considerably low standard deviation (< 0.1 m·s^−1^) (30) and estimation errors as low as 2-3% (31). Therefore, conduction velocity estimates of earlier studies were likely less accurate than those obtained in the present study, not only due to the different recording technique adopted but also to the different algorithms applied.

Individual MUCV values laid within the physiological ranges in all cases (41, 44, 45) and displayed a unimodal distribution (Figure 2 B). Our results are in agreement with previous studies on BB (41, 46) and other muscles (26), which did not observe distinct classes of fibres when estimating their MUCV. Similarly, the distribution of muscle fibre CSA derived from muscle biopsies was continuously distributed, despite fibre typing (Figure 2 A). As suggested by Del Vecchio (26), the continuous distribution of motor unit properties (i.e., MUCV, fibre CSA) may be a prerequisite for a smooth generation of force in the full recruitment range and for energy control. Furthermore, the average conduction velocity (MUCV_AVERAGE_) of each participant, i.e., mean MUCV value from all their active motor units in the ascending phase of the ramp contraction, was in agreement with MUCV values observed during voluntary contractions in the BB muscle by previous studies (41, 47).

Another important factor to consider that relates to the methodology adopted to estimate MUCV and to derive MFD in the current study, is the observed independence of MUCV from the subcutaneous adipose tissue interposed between the muscle belly and the recording site (Figure 2 F). Our results contrast with those of previous studies (48, 49), which reported that muscle fibre conduction velocity (MFCV) estimated from the interference EMG, as the weighted mean of the conduction velocities of the concurrently active motor units was influenced by the subcutaneous fat layer. Whereas in the current study we found that the CV of individual motor units (i.e. MUCV) was not influenced by the subcutaneous fat layer. Thus, the proposed methodology has the potential to reliably estimate MFD in populations with very different distributions of both muscle fibre size and subcutaneous adipose tissue.

Moreover, in agreement with previous reports on BB and other muscles (26, 33, 34, 36, 45, 50), we found a strong association between MUCV and MU RT (R^2^ = 0.78 ± 0.09) (Figure 2 E). This indicated the progressive recruitment of motor units with increasing propagation velocities, and further confirmed that MUCV can be considered as a robust size principle parameter (26, 32, 33).

In this study we found MUCV to be associated with muscle fibre size, whilst being unrelated to fibre type composition. These findings indicate that fibre size, independent from fibre type composition, determines MUCV. A possible explanation that could justify the existence of a biophysical relation between MUCV and muscle fibre size is the lower cytoplasmic resistance of larger muscle fibres, which results in faster propagation of a MUAP along their sarcolemma compared to smaller muscle fibres. The lower cytoplasmic resistance along with the specific electrophysiological features of larger muscle fibres (51–53), which directly influence the excitability of the sarcolemma, seem to justify the faster propagation of MUAP along the sarcolemma of bigger muscle fibres (i.e., fibres with greater diameter or CSA), generally innervated by larger α-motor neurons and hence generally belonging to HT MU, compared to that of smaller muscle fibres, generally innervated by smaller α-motor neurons and hence generally belonging to LT MU. In line with this explanation of a dependency between MUCV and muscle fibre size, the model proposed in the current study demonstrated the possibility to predict the mean MFD and fibre CSA non-invasively from an EMG decomposition-derived neuromuscular parameter with a relatively low margin of error.

There are some limitations within the current study that should be recognized. Firstly, we established a relation between MFD and MUCV, a decomposition-derived parameter which reflects the average propagation velocity of MUAPs along muscle fibres innervated by individual motor neurons. Consequently, this relation could not be assessed on individual muscle fibres because it is technically impossible to estimate conduction velocity and diameter from one and the same fibre *in vivo*. Furthermore, MFD were indirectly derived from fibre CSA measurements by assuming that the muscle fibre is straight and circular, whereas other studies assumed an elliptical fibre shape (54). Moreover, although the adoption of HDsEMG grids ensured a representative spatial sampling of motor unit behavior and properties that allowed to overcome the limitations of intramuscular EMG recordings (8), we have to consider that the biopsy-derived MFD were likely affected by sampling errors, since only a small proportion of muscle fibres were assessed. Therefore, both MFD and MUCV values might have been affected by a sampling error, whose magnitude is difficult to quantify. Lastly, it has to be noted that in order to assess the validity and generalizability of the proposed interface to estimate muscle fibre size, future studies assessing the relations between the estimated and measured fibre size indicators on an independent sample are needed.

Overall, our results demonstrate for the first time the potential of using an EMG decomposition-derived physiological parameter, i.e. MUCV, to reliably estimate muscle fibre size in-vivo in a fully non-invasive way. Here we confirmed that MUCV can be considered not only as a reliable and valid indicator of the progressive recruitment of motor units, i.e., a size-principle parameter, but also as a novel strong biomarker of muscle fibre size. The proposed neuromuscular interface opens new perspectives in the use of MUCV not only in the study of neuromuscular diseases but also in other fields where the non-invasive and painless determination of muscle fibre diameter or cross-sectional area, becomes a priority, such as in aging, space and exercise physiology.

## MATERIALS AND METHODS

### Participants

The participants involved in the present study were the same as in our recent publication (40), which compared motor unit behavior during voluntary contractions in chronically resistance-trained athletes (RT) versus a cohort of healthy untrained individuals (UT). Here we report data from MUCV estimated from HDsEMG and muscle fibre size parameters derived directly from muscle biopsies.

Thirty healthy young males volunteered to take part to this cross-sectional study and provided written informed consent prior to their participation. As general requisites for inclusion, volunteers were required to be aged 18-40 yr, without any underlying health issues, with no history of traumatic upper body injury and/or surgery and with no self-reported history of androgenic-anabolic steroid use. Specific inclusion criteria for the RT cohort were extensive history of upper limb resistance training (≥ 2 sessions·wk^−1^ for ≥ 10 mo·yr^−1^, for ≥ 3 yr) and elbow flexion isometric maximum voluntary torque (iMVT) > 90 Nm. Specific inclusion criteria for the UT group were no history of regular upper limb resistance training and no involvement in systematic physical training in the 18 months prior to their participation in the study. Considering the aim of this current analysis, participants were treated as one single heterogeneous group, presenting a wide variability in terms of elbow flexor maximal voluntary isometric force, muscle fibre size and anatomy (*Table 1*).

**Table 1.** Anthropometric, physical activity, muscle strength, fibre size and muscle-electrode distance of the 29 participants. Data are presented as group mean ± SD.

| Variables | Group<br>(n = 29) |
| --- | --- |
| Age (y) | 21.9 ± 3.2 |
| Height (m) | 1.80 ± 0.08 |
| Body mass (kg) | 82.1 ± 13.9 |
| IPAQ score (MET min·wk<sup>-1</sup>) | 4047 ± 2496<br>range, 825 - 12252 |
| Maximum voluntary force (N) | 376.5 ± 104.1<br>range, 195.2 - 566.6 |
| Muscle fibre CSA (μm<sup>2</sup>) | 6815 ± 2122<br>range, 3006 - 11413 |
| Muscle fibre type I CSA (μm<sup>2</sup>) | 6068 ± 1656<br>range, 2643 - 9332 |
| Muscle fibre type II CSA (μm<sup>2</sup>) | 7524 ± 2658<br>range, 3385 - 13624 |
| Muscle-electrode distance (mm) | 4.4 ± 1.3<br>range, 2.7 - 7.8 |

One participant was excluded from the analyses because the biopsy sample did not yield sufficient useful muscle tissue. Thus, here we present the results for twenty-nine participants (Table 1)

The experiments were carried out at Loughborough University (UK), the protocols and procedures were approved by the Loughborough University Ethics Review Sub-Committee (R17 – P174) and conformed to the requirements of the Declaration of Helsinki.

### Overview

Participants visited the laboratory on three occasions for measurements of the elbow flexors, and specifically biceps brachii muscle, of the non-dominant arm (defined as the non-writing hand). The familiarization session (Visit one) involved habituation with the experimental procedures and the performance of a series of maximum and submaximal isometric contractions with the elbow flexors. Participants’ physical activity was assessed with the International Physical Activity Questionnaire (IPAQ, short format, (55)), and the RT group completed a resistance training questionnaire and follow-up individual discussion. These measurements ensured that only participants that fit the criteria for the two groups (described above) were retained in the study. The main neuromuscular assessment (Visit two), 7-to-10 days after visit one, involved the simultaneous recording of elbow flexors force with isometric dynamometry, and HDsEMG for MUCV estimation from the BB muscle during both maximum voluntary contractions (MVCs) and submaximal force-matching ramp contractions. Prior to HDsEMG, participants underwent B-mode ultrasonography to quantify the muscle-electrode distance (MED), i.e., an indicator of subcutaneous adipose tissue thickness, and its influence on MUCV estimation. Approximately, 7-to-8 days after visit two, participants reported to the laboratory for the muscle biopsy (Visit three), which was taken in the region of HDsEMG recordings, for the assessment of muscle fibre diameter and CSA. Participants were instructed not to participate in demanding physical activity (48 h) and to avoid caffeine (24 h) consumption before each session.

## EXPERIMENTAL PROCEDURES

### Neuromuscular assessment

In visit two, after a standardized warm-up (40), participants performed 3-4 MVCs with the elbow flexors of their non-dominant arm, and were instructed to “pull as hard as possible” for 3-5 s. Rest between contractions was ≥ 30 s, and strong verbal encouragement to overcome the peak force of the previous MVCs was displayed with a horizontal cursor. The highest instantaneous force recorded during any MVC was defined as elbow flexor maximum voluntary isometric force (MVF), and was used as reference to define the submaximal force levels.

Five minutes after the MVCs, participants performed 8 submaximal ramp contractions; two contractions to each force level of 15%, 35%, 50% and 70%MVF in a randomized order. The contractions involved a linear ascending ramp phase increasing force from rest to the specified target force level, a plateau phase of 10 s (for 15-35%MVF trials) or 5 s (for 50-70%MVF trials) of constant force production at the target force level. The rate of force increase was kept constant at 10%MVF·s^−1^ for all contractions, and trials were separated by 3-5 minutes of rest. During the aforementioned tasks, participants were instructed to match as precisely as possible a visual template displayed on a monitor positioned at a 1-m distance from the participants’ eyes, for 5 s before, and throughout each contraction. The real-time force output was superimposed on the template for visual feedback.

### Force recording

Participants neuromuscular force production was recorded whilst they were seated in a custom-built isometric elbow flexion dynamometer, consisting of a rigid strength-testing chair, which was adjusted to the participants’ height and upper arm length. Participants were seated in an upright position (hip joint angle at ~90°) with their non-dominant shoulder at ~90° of flexion (i.e. perpendicular to the trunk) and slightly horizontally abducted at ~10°, with the elbow at ~70° of flexion (0° = full extension), and with forearm in half-supination ~45° (~0° = anatomical position) (40). The wrist was tightly strapped to an adjustable wrist brace in series with a calibrated S-beam strain gauge (Force Logic, Swallowfield, UK), positioned perpendicular to forearm rotation. Participants were firmly strapped to the seat back across the pelvis, chest and shoulder to limit extraneous movements.

The analog force signal was amplified (x200), digitized at 2048 Hz using the same acquisition system as that used for HDsEMG (EMG-Quattrocento, OT Bioelettronica, Turin, Italy), and recorded with the software OT BioLab Ver. 2.0.6352.0 (OT Bioelettronica, Turin, Italy). Real-time force visual feedback as well as the ramp templates were provided with the computer software Spike 2 (CED, Cambridge, UK).

### Ultrasound recording

Prior to electrode placement, ultrasonographic (US) images of BB muscle were captured at rest using B-mode ultrasonography (EUB-8500; Hitachi Medical systems UK Ltd., Northamptonshire, UK) and a 92-mm, 5-10 MHz linear-array transducer (EUP-L53L). Resting US images were captured with the participants standing in an upright position with their non-dominant shoulder at ~90°of abduction (i.e., humerus perpendicular to the trunk), with elbow fully extended and forearm supinated. Participants were instructed to stand still and relax their arm, elbow and shoulder, which were supported by adjustable padding. A surgical pen was used to delineate the profiles of the two BB muscle bellies (i.e., short and long head) firstly identified through palpation, and a central line running along the length of each belly at 50% of the medio-lateral width was drawn. Narrow Echo-absorbent markers were placed on the dermal surface (perpendicular to the length of the humerus) from the muscle-tendon junction of BB (i.e., 0 cm) at 5 cm intervals (i.e., 5, 10, 15 cm) along the length of the muscle bellies. Water soluble transmission gel was then applied above each muscle belly to optimize US image detection along the entire length of the muscle. The transducer, also coated with water soluble transmission gel, was gradually moved with minimal pressure applied on the dermal surface along the center line of each belly (i.e., short head and long head) from the distal BB muscle-tendon junction to the proximal end of the muscle. A single sweep of US images was performed and recorded (see below) for each muscle belly.

US was used to quantify the muscle-electrode distance (MED), which was adopted as an indicator of subcutaneous adipose tissue thickness interposed between the muscle belly and the recording electrodes. The Echo-absorbent markers were adopted to align three separate images (i.e., 0-5 cm, 5-10 cm and 10 cm+). This allowed measurement of MED, i.e., distance from the surface of the skin to the muscle fascia, at 1 cm intervals along the length of BB muscle. Specifically, MED was measured starting at −2 cm distal relative to the placement of the distal edge of the HDsEMG grid, and then every 1 cm until 13 cm above the starting location (HDsEMG grid length of 10.9 cm). This approach was used to at least partly account for distal movement of the underlying muscle beneath the HDsEMG grid, between the US measurement (i.e., elbow fully extended) and elbow flexion contractions (i.e., elbow at ~70° of flexion). Furthermore, in order to assess the influence of MED on MUCV estimation, only the average MED values measured from the short head of BB were correlated with MUCV_AVERAGE_ values recorded from the muscle region underlying the electrode grid.

The video output from the US was transferred to a computer and US images were recorded through Ezcap Video Capture software. US images analyses were performed with the Tracker image analysis software (Ver. 5.0.6 http://www.physlets.org/tracker/).

### HDsEMG recording

Myoelectrical activity was recorded from the BB muscle during the submaximal isometric contractions using two high-density surface adhesive grids with 64 electrodes each (13 rows x 5 columns, 1 mm electrode diameter, 8 mm inter-electrode distance; GR08MM1305, OT Bioelettronica, Turin, Italy; Figure 1 B). This electrode configuration formed an array of 128 electrodes covering the short and long heads of the BB. To ensure an optimal placement and orientation of the electrodes, an experienced investigator identified the two bellies of the BB long and short heads through B-mode ultrasonography, and marked their profiles with a surgical marker. The portion of the skin identified using the criteria defined above was shaved, gently abraded and cleansed with alcohol wipes. The electrode grids were positioned with the participant’s non-dominant arm in the testing configuration (~70° of elbow flexion). The grids were centered between the proximal and distal tendons of the BB muscle (average distance of the distal side of the array from the antecubital fossa, 5.9 ± 1.2 cm), with the electrode columns aligned with the assumed direction of muscle fascicles (34, 45, 56). Disposable bi-adhesive foam layers, whose cavities were filled with conductive paste (SpesMedica, Battipaglia, Italy) to optimize the skin-to-electrode contact, were used to attach the electrode grids to the skin overlying the BB. The large number of electrodes covering the BB muscle bellies allowed the identification of the innervation zone (IZ) in the proximal portion of the muscle, and the visual selection of recording electrodes with propagating MUAP along the entire length of the grids. Reference electrodes were located on the ulna styloid process (i.e., main ground electrode) and the radial styloid process (i.e., high-density grids references).

HDsEMG signals were acquired in monopolar derivation, amplified (x 150), band-pass filtered (10-500 Hz) and digitized at a sampling rate of 2048 Hz using a multichannel 16-bit A/D acquisition system (EMG-Quattrocento, OT Bioelettronica, Turin, Italy). HDsEMG signals were recorded with the software OT BioLab Ver. 2.0.6352.0 (OT Bioelettronica, Turin, Italy) and synchronized with the force signal at source.

### Muscle biopsy

During visit three (8.4 ± 5.4 days from Visit two), muscle biopsies were taken from the BB under local anesthesia (1% lidocaine) using the Well-Blakesley conchotome technique (18). Muscle biopsies were taken from the distal anterior BB (average distance of the biopsy from the antecubital fossa, 6.6 ± 0.9 cm) in the same region the HDsEMG recordings were made. Muscle samples were dissected of any visible connective tissue and fat, and then immediately embedded in a mounting medium (Tissue-Tek O.C.T. Compound, Sakura Finetek Europe, Alphen aan den Rijn, The Netherlands) and frozen in liquid nitrogen-cooled isopentane. Samples were then stored at −80°C for immunohistochemistry analysis.

## DATA ANALYSIS

### Force analysis

The acquired force signal was converted to newtons (N) and low-pass filtered (4^th^ order, zero-lag Butterworth filter, 15 Hz cut-off frequency). The offset was removed by correcting for the effect of gravity. Only one of the two isometric ramp contractions at each force target (15, 35, 50, 70%MVF) was analyzed for each participant. The selection criteria was the lowest deviation of the force trajectory from the given template (36, 40).

### HDsEMG decomposition

Monopolar HDsEMG signals were initially band-pass filtered (2^nd^ order, zero-lag Butterworth filter, 20-500 Hz cut-off frequencies) and visually inspected for noise and artifacts. Channels with noise or artifacts were excluded from further analysis. The filtered signals were decomposed into individual motor unit pulse trains with the Convolution Kernel Compensation decomposition algorithm implemented in the DEMUSE tool software (Ver. 5.01; The University of Maribor, Slovenia) (11, 28). This decomposition approach accurately identifies motor unit discharge times over a broad range of volitional forces and has been extensively validated using experimental (57) or simulated signals (11). After the automatic identification of individual pulse trains, the decomposition accuracy was first assessed by calculating the pulse-to-noise ratio (PNR) for each motor unit. The PNR metric is correlated with both the sensitivity and false alarm rate in the identification of motor unit discharges (11). The identified motor unit pulse trains from both grids of electrodes were visually inspected and manually edited by an experienced investigator to check for false positives and false negatives (58) according to available guidelines (9). Only motor units exhibiting a reliable discharge pattern with a PNR > 30dB (sensitivity > 90%, false alarm rates < 2%) and/or interspike intervals < 2s were retained. Furthermore, only motor units extracted from the grid of electrodes, which showed the highest number of reliably identified motor units, were included in the following analyses. In all cases (29/29), this was the grid of electrodes located on the short head of the BB (40).

For each retained motor unit, basic properties were extracted. Specifically, the normalized recruitment threshold (MU RT) and the discharge rate (MU DR) in the ascending phase of the ramp contractions were computed for all identified motor units. Normalized MU RT was defined as the percentage of force (%MVF) generated by the elbow flexors at which the first MUAP was discharged. MU DR was calculated as the average of the first 20 discharges (MUAPs) of each motor unit during the ascending phase of the ramp. This number of discharge timings was chosen to minimize the effects of interspike interval (ISI) variations both on the computation of MU DR in the ascending phase of the ramp contractions and on the estimation of MUCV (31, 34, 59). For clarity, data across the four different target forces (15, 35, 50, 70% MVF) for each subject were collapsed to produce overall motor unit properties (MU RT, MU DR, MUCV), irrespective of contraction level (40).

### Motor unit conduction velocity estimation

Since the decomposition algorithm identifies only individual pulse trains but not the waveforms of the corresponding multi-channel MUAP, these were obtained via the spike triggered averaging technique (31, 59). The multichannel MUAP waveforms were estimated by averaging the recorded monopolar (raw) EMG signals over 15 ms intervals (MUAP duration), using the first 20 discharge times of each motor unit as triggers (31, 34).

Double differential (DD) derivations were computed by differentiating the averaged monopolar MUAP along the electrode columns of the high-density grids, and were used for MUCV estimation (31). A minimum of 3 up to a maximum of 8 DD channels from the same electrode column were visually inspected and selected for MUCV estimation for each identified motor unit. The visual selection of the highest number of DD EMG channels yielding the clearest propagation of MUAP along the electrode columns with minimal change in MUAP shape (from IZ to the distal tendon region) and with the highest cross-correlation (CC) between consecutive channels (CC ≥ 0.70), is the most accurate approach for EMG channel selection and for MUCV estimation (31, 32, 34). Once the channels were selected, MUCV was estimated through a validated multichannel maximum likelihood algorithm that ensures the accurate calculation of propagation velocity in single motor units with a low standard deviation (< 0.1 m·s^−1^) (30) and estimation errors (2-3%) (31). Firstly, MUCV was estimated for each of the 701 motor units identified from BB muscles of all the participants. Secondly, the average MUCV value of the identified and active motor units in the ascending phase of the ramp contractions was computed (MUCV_AVERAGE_).

### Muscle fibre analysis

Transverse serial cross-sections (8 μm thick) of muscle tissue were obtained using a cryotome and placed onto poly-L-lysine-coated glass slides. Sections were fixed for 10 min in 3.7% paraformaldehyde at room temperature and blocked with tris-buffered saline containing 5% goat serum, 2% bovine serum albumin, and 0.2% Triton for 1 h at room temperature. Serial muscle sections were then incubated with a primary antibody for myosin heavy chain I (A4.951, Developmental Studies Hybridoma Bank, Iowa City, USA) and diluted 1:200 in the blocking solution for 1 h at room temperature. Sections were then incubated for 2 h at room temperature with an appropriate secondary antibody consisting of goat anti-mouse Alexa Fluor 488 (A11029, Fisher Scientific, Pittsburgh, USA) diluted 1:500 and wheat germ agglutinin Alexa Fluor™ 350 Conjugate (W11263, Fisher Scientific, Pittsburgh, USA) diluted 1:20 in the blocking solution. Following incubation, coverslips were mounted with Fluoromount aqueous mounting medium (F-4680, Sigma-Aldrich, St. Louis, USA).

Images were captured at x20 magnification using a fluorescence microscope (Leica DM2500). All fibre image analyses were performed using Fiji (ImageJ) software (60), and the investigator was blinded to the participant group allocation. Only transversely sectioned fibres were included in the analysis (i.e., any fibres that were clearly oblique or not transverse to the long axis of the fibre were excluded). Fibre cross-sectional area (fibre CSA) was assessed by manually drawing around the perimeter of each fibre for ~200 different fibres per participant. In a small number of participants (n = 4), only 145-165 fibres were analyzed for area due to limited number of clear images/fibre perimeters.

### Estimation of muscle fibre size from motor unit conduction velocity

First, by assuming that muscle fibre are of uniform cylindrical shape (i.e., circular in cross-section), we converted the single muscle fibre CSA values (*n* = 5619) obtained from muscle biopsies into measured MFD, as follows:

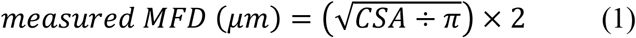

Second, we converted single propagation velocity values estimated from the overall population of identified motor units across all subjects into estimated MFD with Nandedkar and Stålberg’s equation (42), as follows:

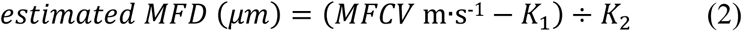

This equation was based on preliminary evidence that reported a biophysical proportionality between the size of a biological conductor, such as a muscle fibre (or axon), and the propagation velocity of action potentials in both animal (37) and human skeletal muscle (61). The two parameters *K*_1_ and *K*_2_ have been estimated in previous studies but in conditions different from the current study (e.g., with intramuscular EMG signals and with electrical stimulation of the fibres). Therefore, they needed to be identified (calibrated) based on the current data.

For calibration, we fitted the estimated MFD data derived from *Equation 2* with the experimentally-derived muscle fibre biopsy data (*Equation 1*) and identified the optimal values of the parameters *K*_1_ and *K*_2_ that minimised the mean square error between the estimated and experimental distributions of MFD. This led to the following equation:

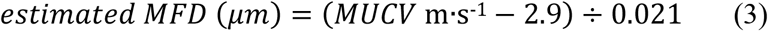

*Equation 3* was derived for the first time from inter-subject observations of both fibre diameters from muscle biopsies and diameters estimated from MUCV. To validate the proposed procedure in terms of predictability, we then adopted a leave-one-out calibration.

The leave-one-out approach consisted in estimating the parameters in *Equation 2* using all dataset except for one subject and repeating the procedure for all subjects. *Equation 2* with the parameter values optimized on all subjects except for one were used to estimate the MFD distribution from the MUCV distribution for the left-out subject. In this way, the estimates of MFD for each subject were obtained without the use of the biopsy data of the subject but rather using a database of MUCV and biopsy data from a separate group of subjects. This analysis provided a direct indication of the quality of estimates of MFD directly from MUCV data. From the leave-one-out estimates of MFD of all subjects, correlation and regression analyses were performed to assess the relation between measured and estimated MFD at the individual subject level.

Lastly, we also explored the relation between muscle fibre CSA estimated from MUCV and muscle fibre CSA directly measured from muscle biopsies, with the same leave-one-out procedure as described above for MFD. Specifically, the estimated MFD obtained from *Equation 3* were converted into estimated CSA as follows:

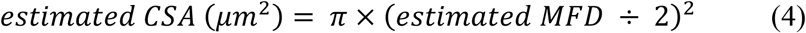

Further correlation and regression analyses were performed to assess the relation between measured and estimated mean CSA.

### Statistical analysis

The Shapiro-Wilk test was used to check the normality of the distribution of data for all the variables analyzed. Corresponding nonparametric tests were used where non-normally distributed data were observed. However, most of the variables analyzed exhibited a normal distribution.

The relative frequency histograms of both individual fibre CSA values (Figure 2 A) and individual MUCV estimates (Figure 2 B) were composed of 41 bins with a ~500 μm^2^ width and 44 bins with ~0.07 m·s^−1^ width.

Independent Student’s *t* test were adopted to assess differences between fibre type I and type II CSA (Figure 2 C) and between MUCV_AVERAGE_ of LT and HT MU (Figure 2 D).

The relation between MUCV and normalized MU RT (%MVF) of all the identified motor units identified for each participant was assessed with Pearson product-moment correlation coefficient (r). The coefficient of determination (R^2^) of the linear regressions was computed for each participant and averaged across participants as an index of prediction power (Figure 2 E).

The relation between MUCV_AVERAGE_ and muscle-electrode distance (MED) was assessed with Pearson product-moment correlation coefficient (r), at the whole population level (Figure 2 F). This analysis was performed to assess the potential effect of subcutaneous adipose tissue thickness on MUCV estimates.

Individual MUCV estimates of all identified motor units were converted into estimated MFD by *Equation 2*. Individual fibre CSA values were converted into measured MFD by *Equation 1*. The relative frequency histograms of both estimated and measured MFD were composed of 170 bins with a ~2 μm diameter increment (Figure 3 A). Similarly, the relative frequency histograms of both estimated and measured fibre CSA were composed of 81 bins with a ~250 μm^2^ area increment (Figure 3 D). The Kolmogorov-Smirnov test was adopted to compare the estimated and the measured MFD/fibre CSA distributions of data.

The relations between estimated mean MFD/mean fibre CSA and measured mean MFD/mean fibre CSA were assessed with Pearson product-moment correlation coefficient (r) at the individual subject level. The coefficient of determination (R^2^) of the linear regressions was computed as an index of prediction power (Figure 3 B, E). The agreement between MFD/fibre CSA estimated from MUCV (estimated mean MFD/fibre CSA) and MFD/fibre CSA derived from muscle biopsies (measured mean MFD/fibre CSA) was assessed with Bland-Altman analysis. The average absolute bias across subjects, i.e., the average of the absolute difference between measured and estimated MFD/fibre CSA for each participant, was determined between estimated mean MFD/fibre CSA and measured mean MFD/fibre CSA. Similarly, the average relative bias across subjects, i.e., the average of the relative difference between measured and estimated MFD/fibre CSA for each participant, was computed.

The absolute and relative errors for each individual were determined by calculating the difference between estimated mean MFD/fibre CSA and measured mean MFD/fibre CSA (normalized to measured mean CSA for relative error), before averaging across participants.

All statistical analyses were performed with the software GraphPad Prism, Ver. 9.2.0 (GraphPad Software, San Diego, California) and MATLAB, Ver. 2021 (MathWorks, Natick, MA). Statistical significance was set at α < 0.05 for all tests. Results are expressed as mean ± SD, unless otherwise indicated.

## Acknowledgements

We thank all participants for their time and efforts in completing the study. Figure 1 A,C were created with BioRender and published with permission.

## Author Contributions

A.C., A.D.V., J.P.F., and D.F. conceived and designed research; A.C., A.D.V., T.G.B., S.M., N.M., and M.B.L. performed experiments; A.C., A.D.V., S.M., T.G.B., and S.N. analyzed data; A.C., A.D.V., T.G.B., S.M., N.R.W.M., F.F., S.N., T.M., A.P., J.P.F. and D.F. interpreted results of experiments; A.C. prepared figures; A.C. drafted manuscript; All authors revised it critically for important intellectual content; All authors have read and approved the final version of manuscript. All persons designated as authors qualify for authorship, and all those who qualify for authorship are listed.

